# A functional landscape of human microproteins

**DOI:** 10.1101/2024.06.10.598216

**Authors:** Mathilde Slivak, Sébastien A. Choteau, Lionel Spinelli, Louis Souville, Philippe Pierre, Anne-Ruxandra Carvunis, Erich Bornberg-Bauer, Andreas Zanzoni, Christine Brun

## Abstract

Short open reading frames (sORFs) are widespread yet often poorly annotated, and an increasing number are known to encode functional sORF-encoded peptides (sPEPs, or microproteins). Here, we use a system approach to determine the functions of sPEPs in three tissues: monocytes, skeletal muscle and brain.

We first predicted interactions between sPEPs and canonical proteins and characterized their interaction interfaces. sPEPs preferentially target specific canonical partners rather than interacting promiscuously, and two functional groups emerge from their interaction features. A small subset (∼2%) exhibits canonical protein-like features, with structured domains interacting with diverse proteins involved in tissue-specific functions. In contrast, most sPEPs contain short linear motifs with extensive phosphorylation potential, suggesting they may be involved in signaling.

We then constructed the first sPEP-containing interactome networks and infer sPEP functions from network topology. Most sPEPs appear to act as regulatory elements modulating diverse cellular processes rather than acting directly as functional components. Knowledge-based validations using characterized sPEPs, show that our framework can capture context-specific functions beyond current annotations. Together with their limited evolutionary constraint, the pervasive regulatory roles of sPEPs suggest that they constitute a reservoir of functional novelty and an evolutionarily flexible layer of cellular regulation.

## Introduction

Open reading frames shorter than 100 codons were initially discarded in most gene annotation programs, since they were thought to have no coding potential^1,2^. More recent studies demonstrated that these sORFs (short ORFs) may often encode functional peptides^3–7^, (sPEPs for sORF-encoded peptides, or microproteins). Although ribosome profiling (Ribo-seq) data have revealed that the human genome codes for three million putative sORFs^8^, their functional relevance remains unclear^9,10^. Indeed, sPEPs were shown to be weakly expressed, prone to degradation and poorly conserved. Moreover, analyzing sPEPs by proteomics, e.g. via mass spectrometry, remains challenging due to their short size, low abundance, atypical sequence features, and frequent enrichment in transmembrane regions^11,12^. Despite these limitations, a growing body of evidence supports the existence of a substantial translated “non-canonical” proteome^1^. Notably, during the course of our study, the TRANSCODE Consortium (https://transcodeconsortium.org/) has generated a standardized catalogue of 7,264 non-canonical ORFs identified by Ribo-seq experiments of length >16 aa (amino acids), initiating with ATG^9^. 25% of these were independently confirmed by mass spectrometry^11^. This catalogue has been extended to contain 28,359 high-confidence non-canonical ORFs, among which 10,127 exhibit translation signature scores comparable to those of canonical protein-coding sequences^13^ (i.e sequences which are well annotated and typically refer to evolutionary well conserved proteins with many homologs in model organisms).

As evidence accumulates that sPEPs are stably expressed, the elucidation of their biological functions is now ongoing. In skeletal muscle, several sPEPs are known to regulate calcium homeostasis^14–18^, play crucial roles in muscle development, maintenance, and regeneration, as well as in lipid and glucose metabolism, skeletal muscle bioenergetics and are involved in cellular stress and muscle-related diseases^17^. Brain sPEPs are potentially involved in neurological diseases^19^ and over 100 human-specific candidates display detectable knockout phenotypes^20^. Functionally characterized examples highlight roles in mitochondrial activity, calcium signaling, and neural development^21–23^. In immune cells, particularly in monocytes, proteogenomic analyses have identified numerous sPEPs, some of which can be presented by MHC-I molecules, suggesting roles in antigen presentation and immune surveillance^24–26^. Overall, many sPEPs display marked tissue specificity, reflecting specialized functions adapted to distinct biological contexts, while recurring functional themes emerge across all tissues as well, notably cell proliferation, signaling, cell growth, death, metabolism, and development^3^, underscoring the contribution of sPEPs to a wide range of cellular processes. However, despite recent large-scale efforts expand functional insights, their characterization remains limited. For example, only 754 human sPEPs, out of several hundreds of thousands sORFs detected^27–29^, are annotated in UniProtKB^30^ with evidence at protein level (“reviewed”). Characterizing and deciphering sPEP function at a large-scale is therefore essential. To address this challenge, we here study the protein-protein interactions (PPIs) of these potential sPEPs with the canonical proteins in different tissue types. Indeed, as PPIs drive biological functions, protein functions can be assigned based on the function of neighbors in the PPI network in a ‘guilt by association’ approach^31,32^. Accordingly, we first predicted the interactions between sPEPs and canonical proteins expressed in the same tissue using *mimicINT*^33^, a computational workflow recently developed for inferring PPIs. Second, the sPEP’s functions have been inferred based on the analysis of the PPI networks. The study notably allowed identifying sPEP’s molecular function through their interaction interfaces, their cellular function through their interaction with protein partners, thereby defining their functional landscape.

## Methods

### Collection of sPEPs identified in three human tissues

The sequences of 664,771 sPEPs identified by ribosome profiling were retrieved in FASTA format from MetamORF^27^ (https://metamorf.hb.univ-amu.fr), a curated repository of short open reading frames (sORFs) identified through experimental and computational approaches. From this complete set, we focused our analysis on sPEPs identified in three tissues of interest: monocytes (10,475 sORFs), brain (55,944 sORFs), and skeletal muscle (1,493 sORFs) (Fig. 1a, Supplementary Fig. 1a).

**Fig. 1:**
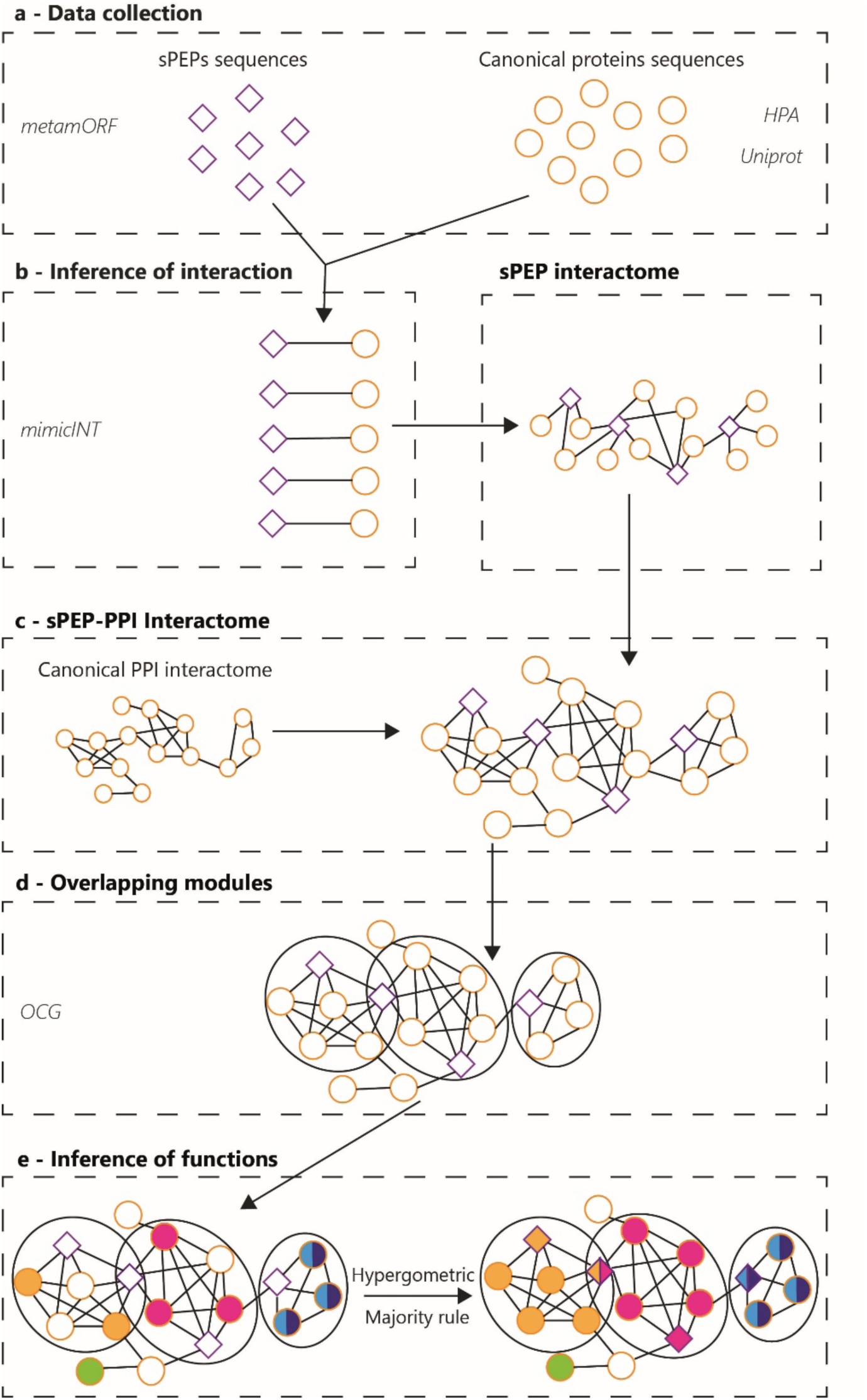
From sPEPs to function. (a) sPEPs sequences identified in monocytes, skeletal muscle and brain have been collected from MetamORF. Canonical protein sequences expressed in each cell type have been collected from UniProtKB and Human Protein Atlas. (b) Binary sPEP-canonical protein interactions are inferred using mimicINT to produce sPEP interactomes. (c) Each sPEP tissue specific interactome is merged with the PPI of the corresponding cell type. (d) The sPEP-PPI interactome is clustered in overlapping modules using OCG. (e) Modules are annotated with GO terms, which are transferred to the sPEPs they contain. Diamond: sPEP. Ellipse: canonical protein.

### Collection of canonical proteins expressed in monocytes, brain and skeletal muscles

Reviewed canonical protein sequences were obtained from UniProtKB^30^ and downloaded in FASTA format (Fig. 1a). Proteins expressed in human monocytes, skeletal muscle, and brain were selected based on experimental evidence of expression reported in the Human Protein Atlas^34^, using a threshold of normalized transcripts per million (nTPM) ≥ 1. (Supplementary Fig. 1a)

### Interaction predictions between sPEPs and canonical proteins with mimicINT

Interactions between sPEPs and canonical proteins expressed in a given tissue type, were inferred using *mimic*INT^33^, a workflow originally developed for microbe–host protein interaction inferences based on templates of Short Linear Motif (SLiM)-domain and domain-domain interactions. Briefly, *mimic*INT first identifies SLiMs on sPEP sequences using SLiMProb^35^, based on SLiM definitions from the ELM database^36^. Detected SLiMs are filtered using *P*-values computed by Monte Carlo simulations (10,000 permutations; p<0.01^37^) to control for false-positives. Globular domains are identified on both sPEP and canonical protein sequences using InterProScan^38^, relying on Pfam signatures^39^. Second, *mimic*INT performs interaction inference by matching detected domains and SLiMs to known templates for domain–domain interactions obtained from 3DID^40^, and for SLiM-domain interaction retrieved from ELM^36^. SLiM-domain interactions were further filtered using a motif-binding domain similarity scores (domain scores) derived from Hidden Markov Model searches^41^ (threshold = 0,4) (Fig 1b).

Interactions involving truncated domains on sPEPs, defined as domains missing more than 10 amino acids relative to the Interpro consensus and therefore likely to impair domain function^42^, were excluded from subsequent analyses (Supplementary Fig. 1b). When sPEPs retrieved from MetamORF has an entry in UniProtKB, the MetamORF identifier was retained for further analysis, and all associated interactions (both experimental and predicted) were reassigned to this unified entry.

### Clustering of the sORFs sequences

To minimize isoform redundancy arising from overlapping sORFs, sPEP amino acid sequences were clustered at 95% sequence identity using CD-HIT^43^. The longest sequence in each cluster was selected as representative of the cluster, which was the only one to be incorporated into the interactome for further network analyses (Supplementary Fig. 1b).

### Merging the sPEP interaction network with the canonical protein-protein interaction networks

The sPEP–canonical protein interaction networks were merged with a canonical protein– protein interaction (PPI) network obtained from MoonDB^44^ (2025 update, **Supplementary Data 1**). The PPI network was restricted to canonical proteins expressed in the corresponding tissue, as defined by the Human Protein Atlas. The resulting three networks are hereafter referred to as the “*merged interactomes”* (Fig. 1c, Supplementary Data 2). The same approach has been applied to the Human Reference Interactome^45^ (HuRI) for validation purposes.

### Network Clustering and Annotation

Network clustering was performed using the OCG algorithm^46^ which decomposes the network in overlapping modules of a maximum size of 200 nodes (Fig 1d). These modules are formed by highly interconnected proteins, which tend to be involved in the same cellular processes and may include protein complexes. Modules significantly enriched in sPEPs were identified using a one-tailed hypergeometric test followed by multiple testing correction (Benjamini–Hochberg corrected P-values < 0.05). Protein modules were functionally annotated using Gene Ontology^47^ (GO) Biological Process terms. Functional enrichment was assessed independently for each module using a one-tailed hypergeometric test to evaluate GO term over-representation relative to the network background, followed by Benjamini– Hochberg multiple testing correction. GO terms with a false discovery rate (FDR) < 1 × 10⁻⁵ were considered significantly enriched. Modules that did not reach statistical significance were annotated using a majority rule, whereby GO terms annotating at least 50% of the annotated canonical proteins in the module were retained^31,32^. sPEPs then inherited the functional annotations of the module they belong to (Fig 1e). Modules smaller than 5 nodes and overly generic GO terms were excluded from the analysis. Redundancy among inferred GO terms was reduced using the R package rrvigo^48^ with a similarity threshold of 0.7 .This list of GO terms was then manually simplified using the QuickGO browser^49^. Networks were visualized with Cytoscape^50^ and ClustnSee3.0^51^. Graph-based visualization of inferred GO terms was extracted from the Revigo^52^ web server with a similarity threshold of 0.9.

### Functional enrichment analyses

Custom GMT files including sPEP annotations were generated by retrieving GO annotations for canonical proteins via the QuickGO API^53^ and formatting them using the GMT helper tool using the go-basic^47^ and the Alliance of Genome Resources^54^ subset obo files from Gene Ontology.

Over- and under-representation of GO terms were performed separately for sPEPs and for canonical proteins interacting with sPEPs using g:Profiler Python package.^55^. Multiple testing correction was applied using the Benjamini–Hochberg method, and terms with False Discovery Rate (FDR) < 0.05 were considered significant.

### C-terminal Tail Hydrophobicity

C-terminal tail hydrophobicity (CTTH) scores^56^ were calculated as a proxy for proteasome– ubiquitin–mediated degradation. For each sPEP, the average hydrophobicity was computed using the Kyte–Doolittle scale, in which hydrophobic residues have positive values.

Calculations were performed using the GRAVY portal (www.gravy-calculator.de). For sPEPs longer than 30 amino acids, only the C-terminal 30 amino acids were considered. For sPEPs shorter than 30 amino acids, the full sequence was used to compute the CTTH score.

### Prediction of Chaperone-Mediated Autophagy Motifs

Potential targeting of sPEPs by chaperone-mediated autophagy (CMA) and endosomal microautophagy (eMI) was evaluated by identifying KFERQ-like pentapeptide motifs^57^. Motif detection was performed using the *“motifs from sequence”* option of the KFERQfinder portal (https://rshine.einsteinmed.edu), applying default parameters to identify canonical motifs as well as phosphorylation- and acetylation-activated KFERQ-like motifs.

### Identification of Terminal Intrinsically Disordered Regions

Intrinsic disorder at the N- and C-termini of sPEPs was predicted using IUPred2A^58^ (long disorder option). Terminal regions were defined as the first and last 22 amino acids of each sequence, in accordance with previously established criteria^59^. Terminal intrinsically disordered regions (IDRs) were identified when the predicted disordered segment extended beyond the defined terminal region.

### Identification of Degron Motifs

Short linear degradation motifs (degrons), including N-terminal and C-terminal degrons as well as internal degrons, were identified using the *mimic*INT workflow.

## Results and Discussion

### 1 Inference of the sPEP-canonical protein interactomes in three different tissues

We generated the first large-scale networks of sPEP–protein interactions across three human tissues: monocytes, skeletal muscle, and brain (chosen for the existing knowledge of the functions of some of their sPEPs^17,25,60^). Sixty-four thousands three hundreds ninety-four putative sORF-encoded peptides (sPEPs) identified by ribosome profiling in each tissue type were collected from MetamORF^27^ (10,475 in monocytes, 1,493 sORFs in skeletal muscle and 55,944 in brain). In parallel, the canonical proteins expressed in each of those tissue were retrieved based on expression evidence from the Human Protein Atlas^34^ (Fig. 1a, Supplementary Fig. 1a).

Interactions between sPEPs and canonical proteins were predicted using *mimic*INT, a sequence-based framework that uses templates of interactions between either globular domains, or short linear motifs and domains, and that reaches an experimental validation rate of 70%^61^ (see Methods; Fig 1.b). After filtering out interactions involving truncated domains and clustering overlapping sPEP sequences to reduce isoform redundancy (see Methods), we retained 82,714, 16,080, and 772,344 sPEP–canonical protein interactions, corresponding to 50,010, 7,150 and 409,325 binary interactions between proteins (Table 1) in monocytes, skeletal muscle, and brain, respectively. Most of these interactions were tissue-type specific. They involved 236 sPEPs in skeletal muscle, 1,810 in monocytes and 8,014 in brain, interacting with 886, 1,603 and 8,086 canonical proteins, respectively (Supplementary Fig. 1b).

**Table 1.**
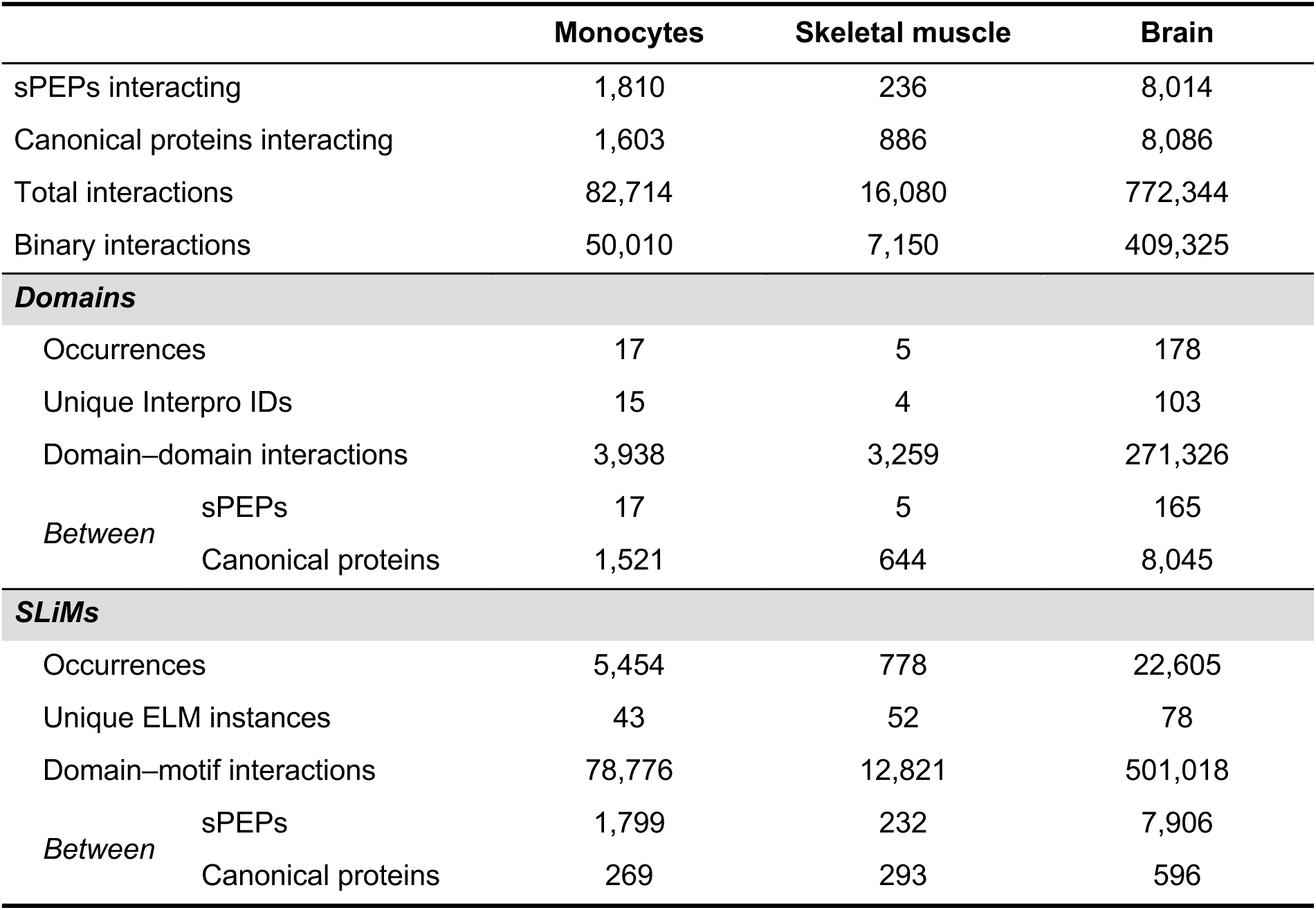
sPEP–canonical protein interactions and domain/motif-mediated interaction statistics across three tissue/cell types.

sPEPs have been proposed to be degraded after translation^62^. We then calculated their propensity to be degraded and showed that none of the tested degradation determinants (C-terminal tail hydrophobicity scores, terminal degrons, terminal intrinsic disorder regions, and the microautophagy-associated KFERQ-like motifs^62^) indicate they are particularly unstable (Supplementary Fig. 2), therefore reinforcing the plausibility of our inferred interactions In each tissue, sPEPs were predicted to interact with a limited subset of the canonical proteome (Table 1). In monocytes, 40% of the sPEPs were predicted to interact with 14% of the canonical proteins, 40% of sPEPs with 7% in skeletal muscle and 36% of the sPEPs with 46% of the canonical proteins in brain (Table 1).

Only 0.07% (7/9710) of the interacting sPEPs were common to the three tissues, and 3% (336/9710) to any two of the three (Supplementary Fig. 1e). In contrast, canonical proteins were more often shared across tissue: whereas 5% (419/8213) were predicted to interact with sPEPs in all three tissues, 19% (1523/8213) do so in two of them (Supplementary Fig. 1f). This indicates that while individual sPEPs tend to operate in a highly context-dependent manner (82 to 95,7% of the sPEPs are specific to only one tissue), canonical proteins are more widely shared as interaction partners across cellular environments.

Despite encompassing tens to hundreds of thousands of predicted interactions per tissue type, sPEP-protein interactions engage only a fraction of the canonical proteome (7 to 46%), indicating that sPEPs do not interact promiscuously but rather target specific canonical interactors. These results suggest that sPEPs contribute significantly to the cellular PPI and act in a highly context-dependent manner on a more broadly shared set of canonical protein partners.

### 2 sPEP SLiMs and domains reveal two types of sPEP functions

We aimed to explore sPEP functions by studying their interactions across multiple contexts. First, interfaces mediating interactions (domains and SLiMs) provide information about the molecular functions of the proteins that harbor them. Indeed, interfaces may mediate interactions with other proteins, notably allowing them to participate in complexes or pathways, to be targeted to certain subcellular compartments, or to undergo post-translational modifications. Hence, we investigated the most used interaction interfaces on sPEPs to get a hint at their putative molecular functions.

As shown in Fig. 2, sPEP–protein interactions were massively mediated by SLiMs rather than domains in all three tissues. Indeed, SLiM–domain interactions accounted for the vast majority of predicted interactions; 95% in monocytes (78,776/82,714), 81% in skeletal muscle (13,523/16,782), and 65% in the brain (501,018/772,344). In monocytes, we identified 5471 interaction interfaces on sPEPs, among which 5454 corresponded to SLiMs and only 17 to domains. Similarly, skeletal muscle sPEPs harbored 778 and 5 interacting SLiMs and domains, respectively, while brain sPEPs contained 22,605 interacting SLiMs and 178 domains (Fig. 2). As expected, these differences in numbers are related to the size of the sPEPs, with the shorter interacting through SLiMs whereas the longer rather use domains (Supplementary Fig. 3). Finally, nearly all sPEPs (98-99%) were predicted to harbor at least one SLiM capable of interactions (Table 1).

**Fig. 2:**
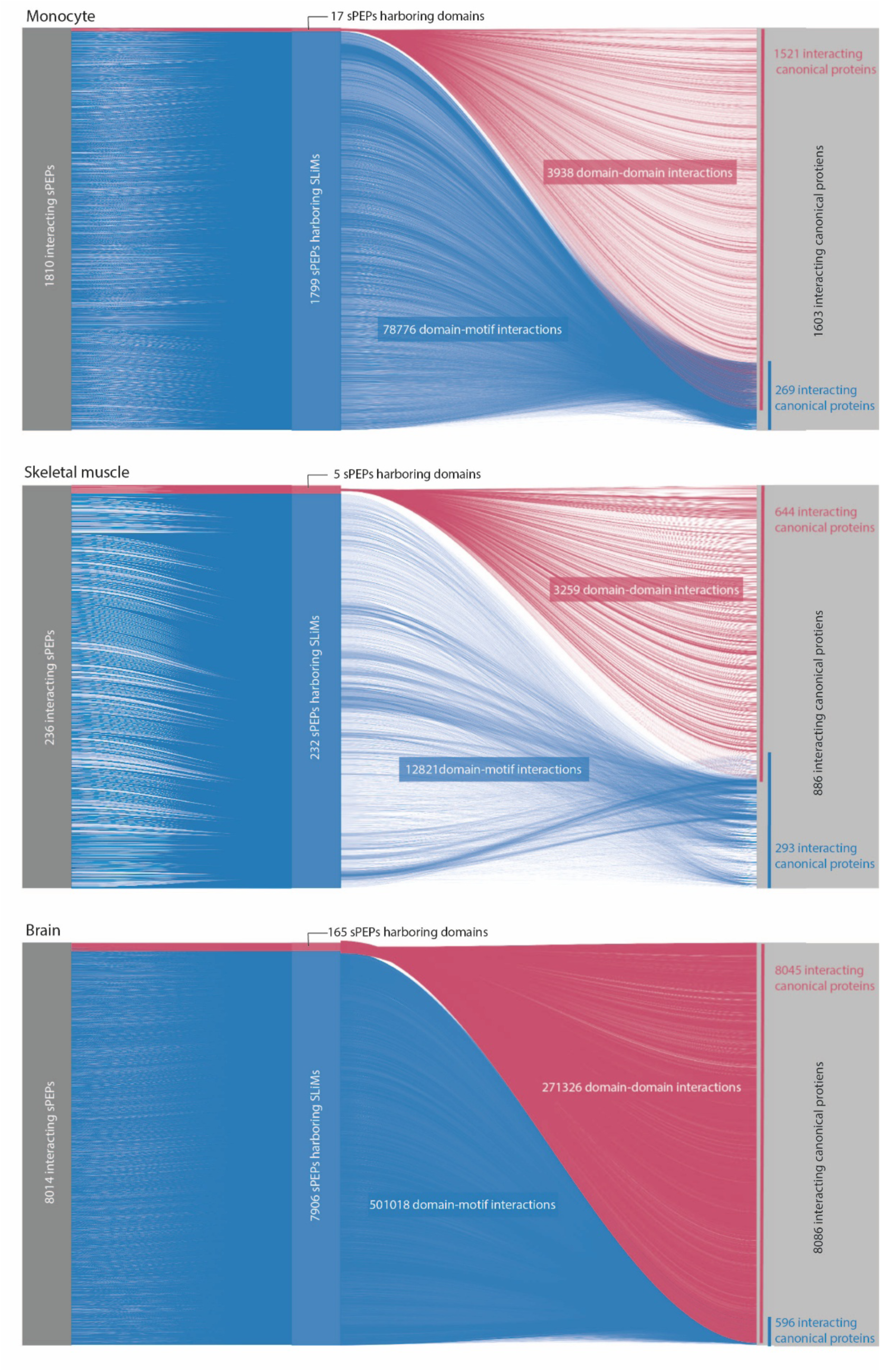
sPEP-canonical protein interactions are mainly mediated by SLiMs. Alluvial plots representing stacked sPEPs of the 3 tissue/cell types (dark grey stack, left side), linked to their interaction interfaces (SLiMs and domains, blue and pink stack, middle), that allow them to interact with canonical proteins (light grey stack, right side).

Among the six SLiM classes defined in the ELM database^36^ — ligand-binding sites (LIG), docking sites (DOC), subcellular targeting sites (TRG), post-translational modification sites (MOD), proteolytic cleavage sites (CLV), and degradation sites (DEG) — MOD motifs were preferentially used by sPEPs to interact with canonical proteins, whereas DEG motifs were the least frequent (4-5% in all tissues) (Fig.3). Strikingly, MOD and CLV motifs are far less numerous among the SLIM’s canonical proteins (∼20% and ∼3% respectively, in all tissues) than among the sPEP ones (between 28-48% and 12-21% respectively, in all tissues). By contrast, canonical proteins preferentially use LIG motifs as SLiM interface (∼50%) (Fig. 3). These differences in SLiM usage clearly indicate that PPIs play distinct molecular roles for sPEPs compared to canonical proteins. Whereas canonical proteins mostly interact with their partners using LIG SLiMs, expected to be mainly involved in stable/permanent interactions, sPEPs favor transient interactions such as phosphorylation or cleavage.

**Fig. 3:**
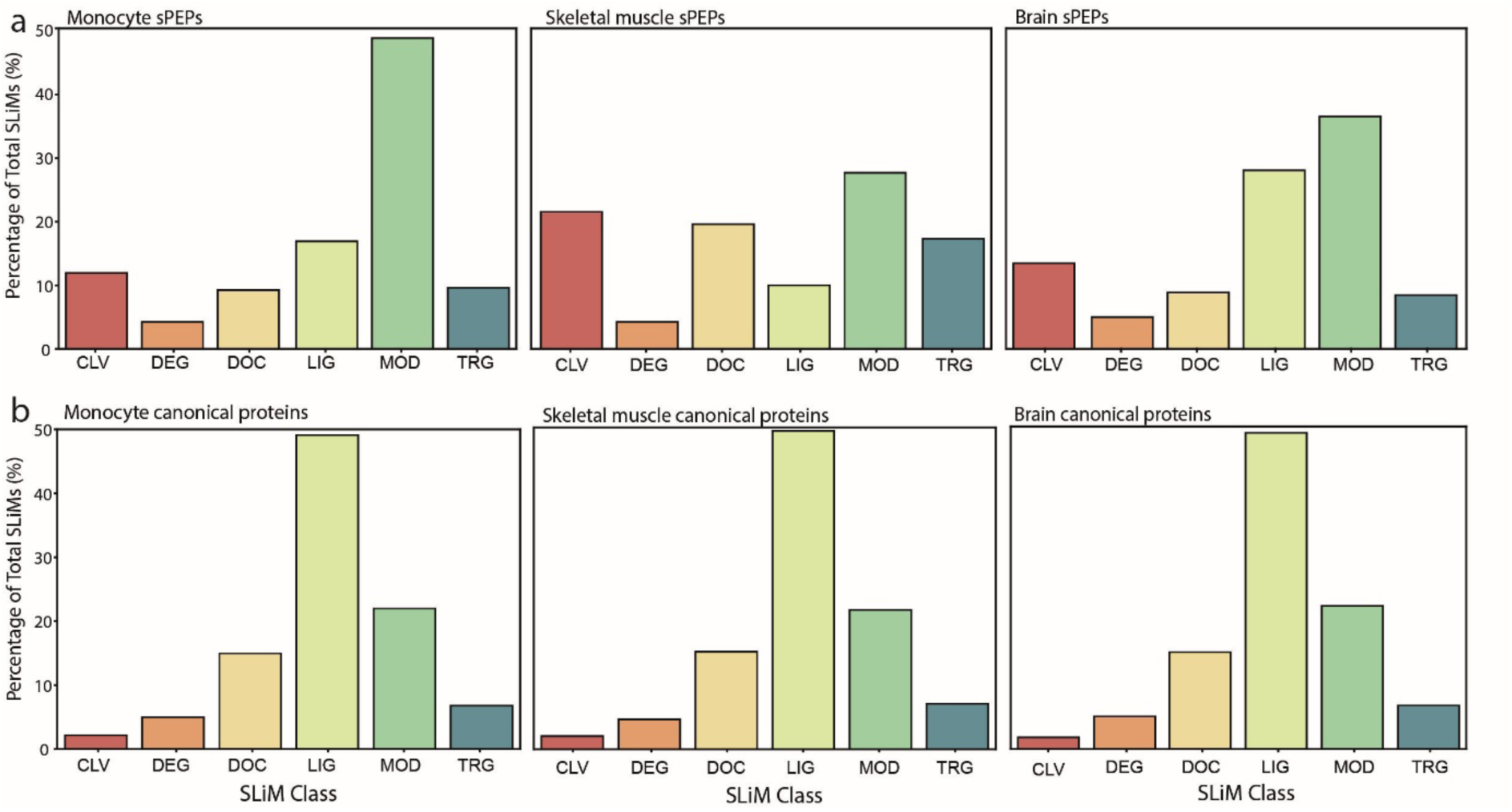
SLiM classes in sPEPs and canonical proteins. (a) SLiM classes predicted in the sPEPs of each tissue type. (b) SLiM classes identified in canonical proteins (= true positive in ELM). CLV = cleavage site, DEG = degron, DOC = docking site, LIG = ligand-binding site, MOD = post-translational modification site, TRG = targeting site.

Although there is little overlap in the sPEP repertoire across tissues, many of the predicted SLiMs are shared: 51% (48/95) of SLiMs are present in at least two tissue types, and 32% (30/95) are found in all three (Supplementary Fig. 1c). Among these, MOD-class motifs mainly interact with the protein kinase domain (IPR000719) in all tissues, suggesting that phosphorylation-related interactions could be a shared feature of sPEPs regardless of their cellular context. Cleavage motifs (CLV) are mainly represented by CLV_PCSK_SKI1_1 and CLV_PCSK_PC1ET2_1 that consistently interact with the same peptidase domain (IPR000209, Peptidase S8/S53 domain) in different canonical proteins and are associated with the proteolytic processing of peptide precursors (Supplementary Fig. 4).

Altogether, these results suggest that sPEPs containing MOD or CLV SLiMs are not only prone to regulation but also regulate some processes upon being phosphorylated or cleaved.

Domains represent only a minor fraction of interaction interfaces on sPEPs across all three tissue types, due to their size (Supplementary Fig. 3). Indeed, domain-mediated interactions involve very few sPEPs (1-2%), and rely on a limited set of InterPro domains (119/8149 possible interaction domains). Accordingly, domain–domain interactions account for a minority of the overall interaction space, representing 5% of interactions in monocytes (3,938/82,714), 19% in skeletal muscle (3,259/16,782), and 35% in the brain (271,326/772,344) (Table 1).

Importantly, despite their low number, domain-mediated interactions connect sPEPs to a large fraction of the canonical proteome in each tissue. In monocytes, skeletal muscle and brain, domain–domain interactions involved 95% (1,521/1,603), 72% (644/886) and 99% (8,045/8,086) of the total interacting canonical proteins respectively (Table 1), in contrast with SLiM-mediated interactions, which involve many sPEPs connected to a lower number of canonical proteins (Fig. 2).

The functional annotations associated with the domains on sPEPs reveal their tissue specificity, *e.g.* small cytokine (intecrine/chemokine) and MHC I C-terminus–associated domains in monocytes, sarcolipin domain in skeletal muscle, and leucine-rich glioma-inactivated (EPTP repeat) and S100/CaBP-9k calcium-binding subdomains in brain.

Remarkably, the coherence of these annotations with the investigated tissue types reinforces the plausibility of the translation of the studied sPEPs. In addition, the fact that only three domains are shared between skeletal muscle and brain sPEPs, none with monocytes, confirms this tissue specificity (Supplementary Fig. 1d).

Overall, the analysis of interaction interfaces supports the idea that first, sPEPs can be largely regulated in all tissues by phosphorylation events and proteolytic cleavages. Notably, the widespread potential of the sPEPs for phosphorylation through SLiMs suggests a possible shared role of the sPEPs as participants or targets of signaling pathways, independently of the tissular contexts. Second, the interaction domains which sPEPs contain, inform on their tissular specificity. Two types of functions, a general and a specific one, can therefore be categorized among sPEPs according to the interaction interfaces they bear.

### 3 The interacting partners of sPEPs inform on sPEP tissular functions

To better understand the functional context of sPEP interactions, we analyzed the functional enrichments in the GO^47^ Biological Process (GO:BP) ^63^ annotations of canonical proteins predicted to interact with at least one sPEP (Supplementary Fig. 5). In total, approximately 25% of the GO:BP terms were shared across all three tissues, with an additional 27% of GO terms common to two tissues, hence underlining a substantial overlap in enriched functions. These were dominated by signaling-related functions: “cell communication” (3.2× 10⁻¹⁵¹< *P*-value < 9 x 10⁻⁵⁰), “signal transduction” (2.6 x 10⁻⁸⁷ < *P*-value < 8.3x 10⁻³⁶), “ phosphorylation” (2.2 x 10⁻¹⁴⁸ < *P-*value < 6.4 × 10⁻³⁹), and “response to stimulus” (6.8 x 10⁻¹²¹ < *P-*value < 1 x 10⁻³³), suggesting that sPEP interactors are consistently involved in signaling pathways. In parallel, each tissue also showed a set of enrichment indicative of its specialized functions, e.g. immune-related processes such as “cell killing” (*P-value* = 4.7 × 10⁻⁸), “antigen processing and presentation of exogenous antigen” (*P-value* = 6.2 × 10⁻⁷), “regulation of leukocyte mediated cytotoxicity” (*P-value* = 7.2 × 10⁻⁹); “muscle hypertrophy” (*P*-value = 1.5 × 10⁻²), “striated muscle hypertrophy” (*P*-value = 1.3 × 10⁻²), “voltage-gated sodium channel activity involved in cardiac muscle cell action potential” (*P*-value = 2 × 10⁻³), “T-tubule organization” (*P*-value = 2.7 × 10⁻²), and “response to muscle stretch” (*P*-value = 4.8 × 10⁻²) in skeletal muscle and specific terms associated to sensory perception such as “detection of chemical stimulus involved in sensory perception of smell” (*P-value* = 5.5 × 10⁻⁴²), and neuronal function such as “regulation of synapse structure or activity” (*P-value* = 9.5 × 10⁻²²), “neuron projection guidance” (*P* = 1.9 × 10⁻¹³) and “trans-synaptic signaling” (*P-value* = 2.1 × 10⁻¹²) in brain.

Overall, on the one hand, common functions of the sPEP-interacting canonical proteins are mainly associated with signaling process functions. On the other hand, tissue-specific mechanisms of sPEPs are reflected by the annotations of their specific interacting proteins, confirming the above analyses on the sPEP interfaces.

### 4 Inference of sPEP functions

Network analyses allow getting an integrated vision of the cellular processes at work in a particular context by notably, focusing on their crosstalks. For this reason, network topology has been successfully exploited in the past to assign cellular functions to uncharacterized proteins^31,32^. Hence, as proteins involved in the same complexes, pathways, or biological processes cluster in the known PPI networks of the canonical proteins, we predicted sPEP functions by analyzing the human contextualized PPI networks to which we merged the sPEP-protein interactions (Table 2). We identified modules in the merged networks using the OCG community clustering algorithm^46^ and annotated them with GO:BP terms (see Methods; Fig 1d-e).

**Table 2:** Overview of the PPI, sPEP, and sPEP–PPI interactomes across three tissue/cell types.

|  | Monocytes | Skeletal muscle | Brain |
| --- | --- | --- | --- |
| **PPI interactome** |  |  |  |
| Proteins | 9,348 | 9,849 | 14,978 |
| Interactions | 57,511 | 63,543 | 94,077 |
| **sPEPs interactome** |  |  |  |
| sPEPs | 1,810 | 236 | 8,014 |
| Interactions | 50,010 | 7,150 | 409,325 |
| **sPEP–PPI interactome** |  |  |  |
| Proteins | 11,158 | 10,085 | 22,895 |
| Interactions | 107,514 | 70,680 | 503,395 |

Among the 375, 322 and 627 modules containing 5 to 200 proteins in the monocyte, brain and skeletal muscle networks, 53%, 97%, and 19% contain at least one sPEP, respectively. Among those, 4 to 19% are statistically enriched in sPEPs (Supplementary Fig. 6). The functional annotations of the modules are then transferred to the sPEPs they contain. By doing so, we predicted the function(s) of 91 to 95% of the sPEPs in monocytes (1644/1810), skeletal muscle (224/236), and brain (7345/8014) networks. sPEPs (Supplementary Fig. 1g).

As sPEP functions are defined with respect to their module’s annotations, they are multi-annotated (10 to 16 terms/sPEP on average). The accuracy of the functional predictions is dependent on the quality of these annotations. To assess the robustness of the functional inferences, canonical protein annotations in the monocyte dataset were randomized while preserving the number of annotations assigned to each protein and the frequency of GO term annotations. Functional inference was then repeated using these permuted annotations, and the results were compared with those derived from the original, non-randomized annotation set. Over 1000 trials, on average, only 5,9% of the sPEPs recover 50 to 100% of their real annotation terms, while 86% of the sPEPs are annotated with less then 10% of their real annotation terms (80% recover 0%). Moreover, the GO terms recovered under randomization were preferentially less specific, with recovery frequency decreasing with increasing GO level and ontology depth (Spearman ρ = −0.26 and −0.48, respectively; P = 3.8 × 10⁻⁶ and 7.8 × 10⁻²⁰). Thus, random annotation assignment rarely reproduced the functional annotation profiles of the sPEPs and, when overlap occurred, preferentially recovered broad rather than specific functional terms. Together, these results indicate that the observed sPEP functional annotations are not explained by random annotation assignment.

Secondly, the functional inferences of the sPEPs were reiterated on the sPEP-protein interactomes merged with the bias-free Human Reference Interactome^45^ (HuRI). Globally, most of the sPEPs are annotated with the same terms as with this HuRI PPI network: while 88%-91% and 70%-95% of the sPEPs are annotated to terms related to “biological regulation” and “metabolic process” respectively, the inferences with HuRI allows predicting 65% to 89% of the sPEPs involved in “biological regulation” and 54% to 91% in “metabolic process”, therefore showing the same trend as with the canonical protein-protein interactome (see above). The preservation of these functional trends using an independent bias-free interactome indicates that the predictions are not strongly dependent on the PPI network used.

Overall, the concordance of these independent assessments (low recovery under randomized annotations, preferential recovery of broad GO terms under randomization, and preservation of functional trends using the bias-free HuRI interactome) reinforces the robustness and biological significance of the functional predictions.

### 5 The functional landscape of sPEPs reveals their involvement in regulatory processes

Studying the functional landscape of sPEPs formed by all the GO:BP terms annotating them, we show that in all tissue types, sPEPs are mainly involved in response to stimulus and metabolic process (Fig. 4). Localization, cellular component organization or biogenesis, immune system process and developmental process are also represented in the functional landscape, although not over-represented (see below). Importantly, for all the processes in which sPEPs are potentially involved, terms related to the regulation of these latter processes are found (Fig. 4). Strikingly, most of the GO terms not related to these processes are, however, related to the regulation of other processes, therefore supporting the relevance of the involvement of sPEPs in regulation at large.

**Fig. 4:**
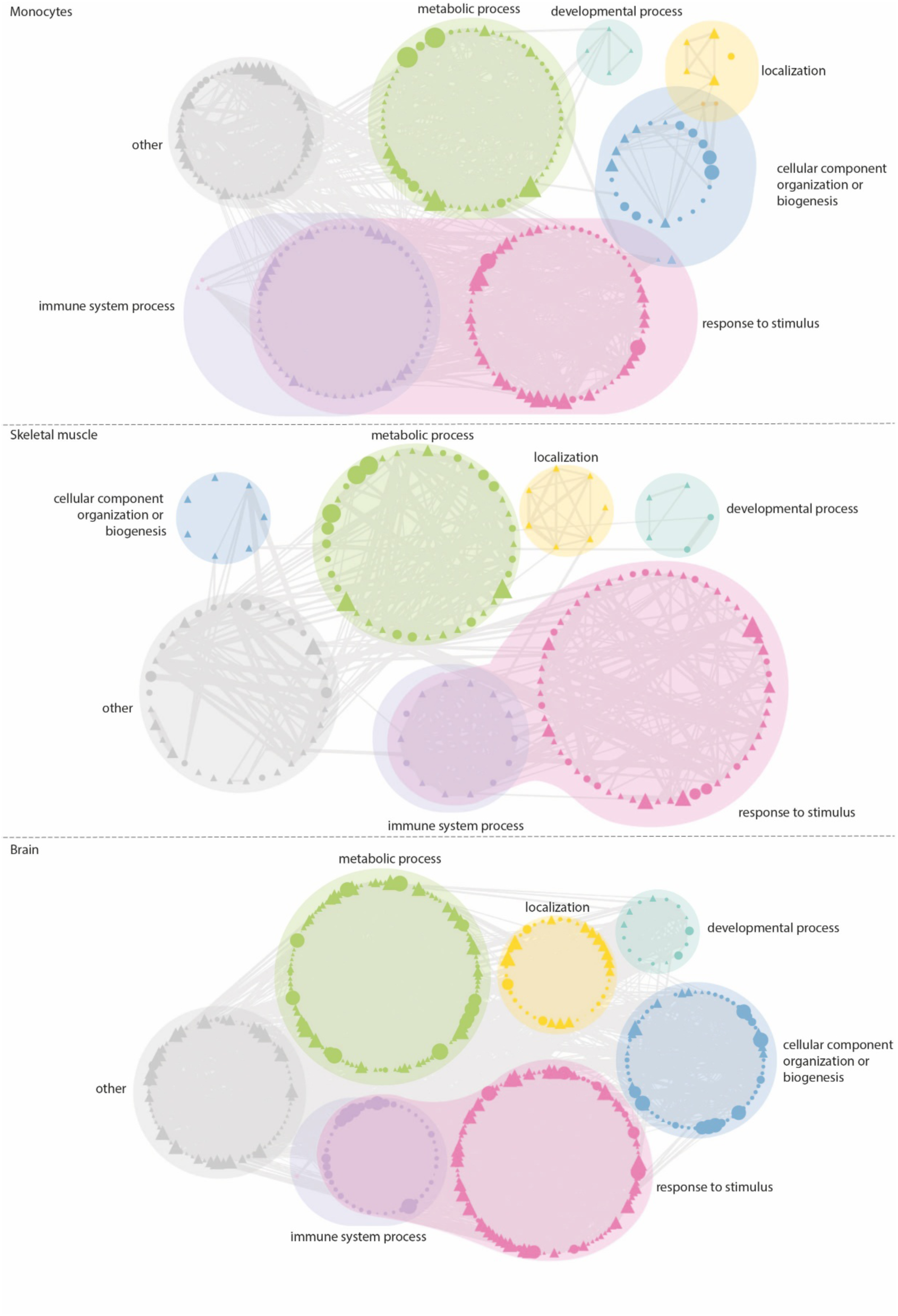
The functional landscape: Nodes represent GO terms annotating the sPEPs, grouped by functions. For each tissue, the size of the nodes is relative to the number of sPEPs annotated to the term in the given tissue. Triangular nodes indicate GO terms related to “regulation”. Light grey edges in the background represent GO links between terms.

Among all the processes present in the functional landscape, enrichment analyses show that sPEPs are consistently highly enriched in functions related to protein metabolic process, cell cycle, signaling and biological regulation (Fig. 5). Noticeably, 31%-44% of the enriched terms are related to regulation in the three tissue types.

**Fig. 5:**
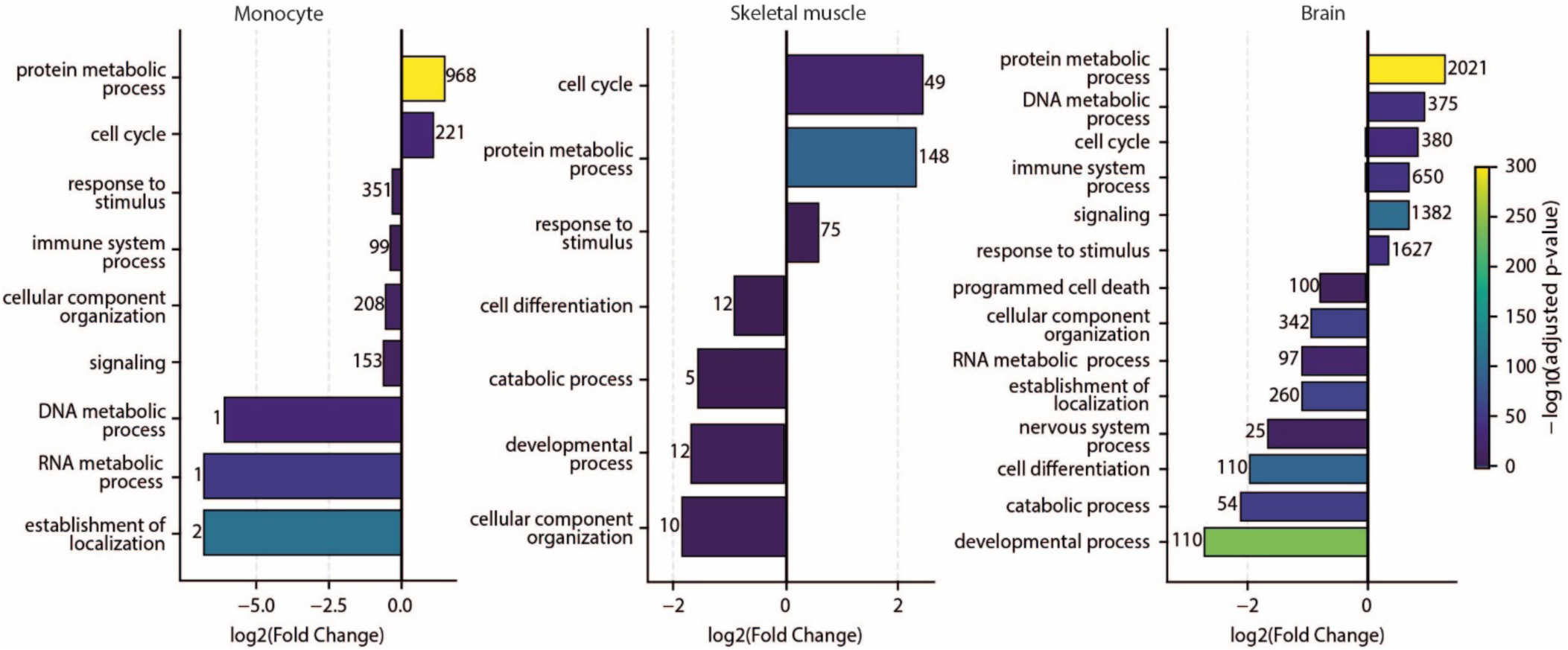
Significantly enriched vs. depleted GO SLIMs among sPEPs. Significantly enriched (positive log₂ fold change) and depleted (negative log₂ fold change) GO SLIM terms among sPEPs in monocytes, skeletal muscle, and brain.

In contrast, several functional categories were consistently statistically under-represented among sPEPs, including “establishment of localization”, “developmental processes” and “cellular component organization”. This under-representation was particularly pronounced in brain and monocytes (Fig. 5). However, numerous terms related to the regulation of these under-represented processes were significantly enriched. For instance, in monocytes, strong enrichments were observed for “negative regulation of inflammatory response”, “regulation of cell communication”, and “regulation of signaling” (*P*-value < 2.5 × 10⁻²²) whereas the processes *per se* are depleted (*P*-value < 3.7 × 10⁻¹⁹). In brain, enriched terms included “regulation of cellular process”, “regulation of biosynthetic process”, “positive regulation of RNA metabolic process” and “regulation of nucleobase-containing compound metabolic process” (*P*-value < 1.9 × 10⁻¹²³), while the non-regulatory terms are highly depleted (*P*-value < 1.6 × 10⁻⁵⁶) (Table 3). These examples highlight the trend of the sPEPs to be consistently involved in regulatory processes across tissues, without necessarily being involved in the regulated functions. This is underlined by the fact that 58 to 97% of the sPEPs of all tissue types are annotated with at least one term related to regulation. Overall, these results therefore confirm the regulatory role of sPEPs for a large number of cellular processes.

**Table 3.**
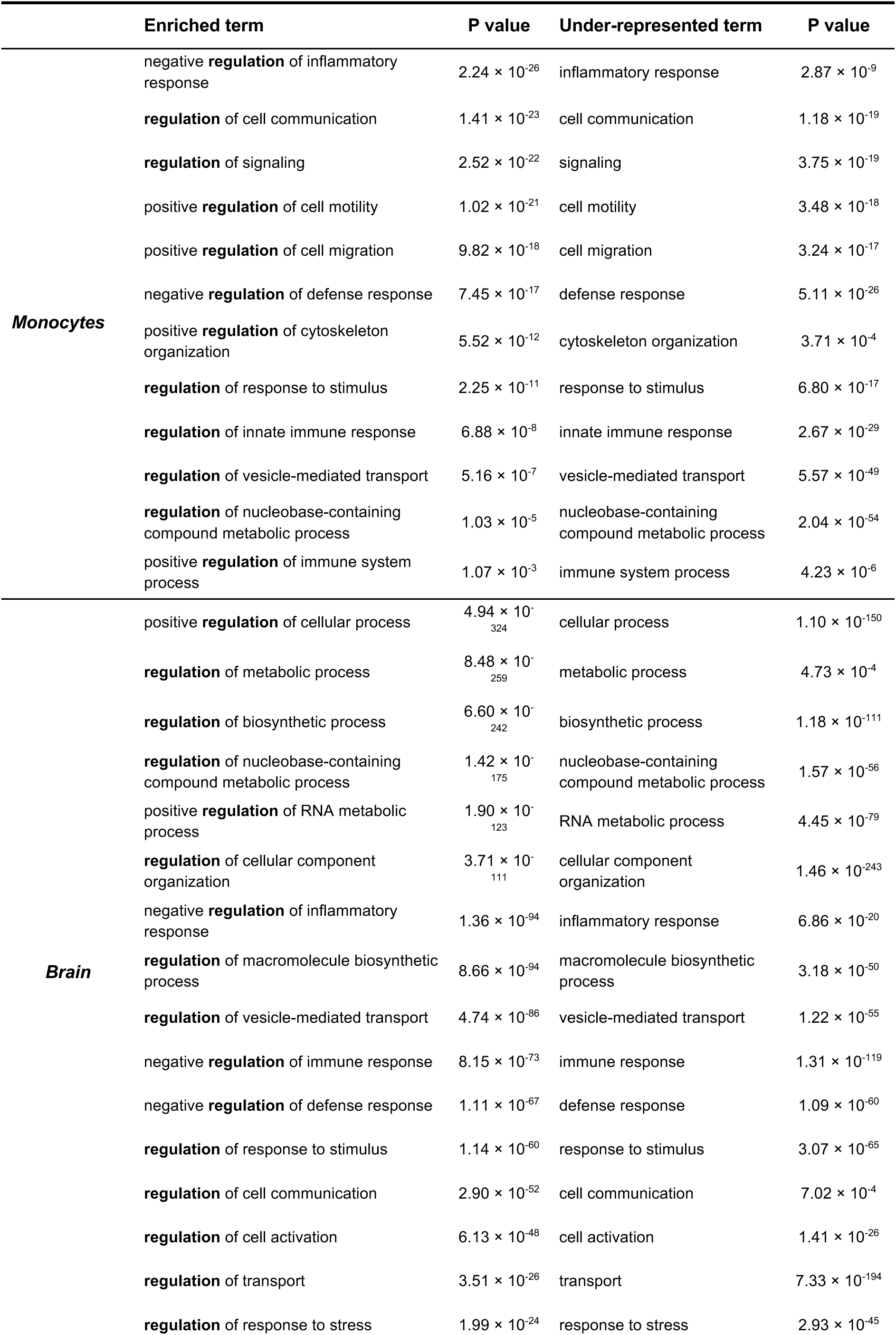
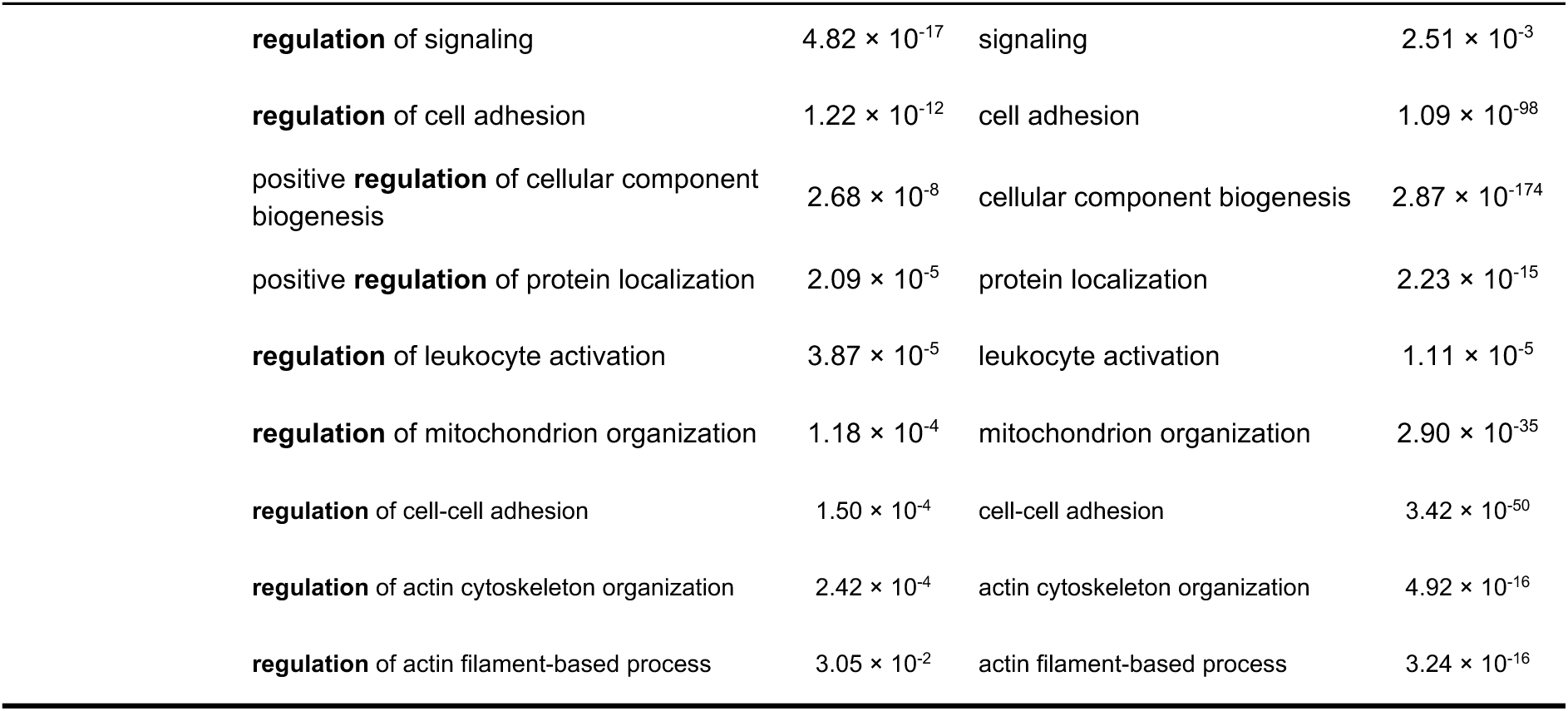
Regulation-related GO:BP terms are enriched among sPEPs in monocytes and brain.

### 6 Case studies of previously annotated sPEPs confirm the inferred functional annotations of sPEPs

Among the sPEPs included in the studied interactomes, the cellular functions of only a few are experimentally demonstrated. We evaluate the annotations assigned to the sPEPs by using sPEPs with known functional roles.

*OST4 in monocytes:* Among the 1,810 sPEPs included in the monocyte interactome, only one corresponds to an annotated protein present in UniProtKB, OST4. Current UniProtKB annotations describe OST4 primarily by its molecular role in “protein N-linked glycosylation”^64^. Consistent with its established function, we inferred direct interactions between OST4 and the catalytic oligosaccharyltransferase (OST) subunits STT3A and STT3B, both supported by domain–domain interaction predictions (Fig. 6a). OST4 has been shown to contribute to the stability and activity of STT3A-containing OST complexes, highlighting its essential role in efficient glycoprotein maturation. As an integral component of the OST complex, OST4 belongs to a functional module annotated to “biological regulation”, suggesting that it indirectly contributes to the regulation of a wide range of biological processes by enabling the glycosylation of numerous proteins with diverse cellular roles (*e.g.*, ER calcium homeostasis and regulation of the ER stress response though glycosylation of WFS1^65,66^). In addition, 15 additional sPEPs were predicted to interact with the catalytic OST subunit STT3B through N-glycosylation sites within this functional module. This interestingly suggests that N-glycosylation may represent, next to phosphorylation and cleavage, yet another common post-translational modification of sPEPs.

**Fig. 6:**
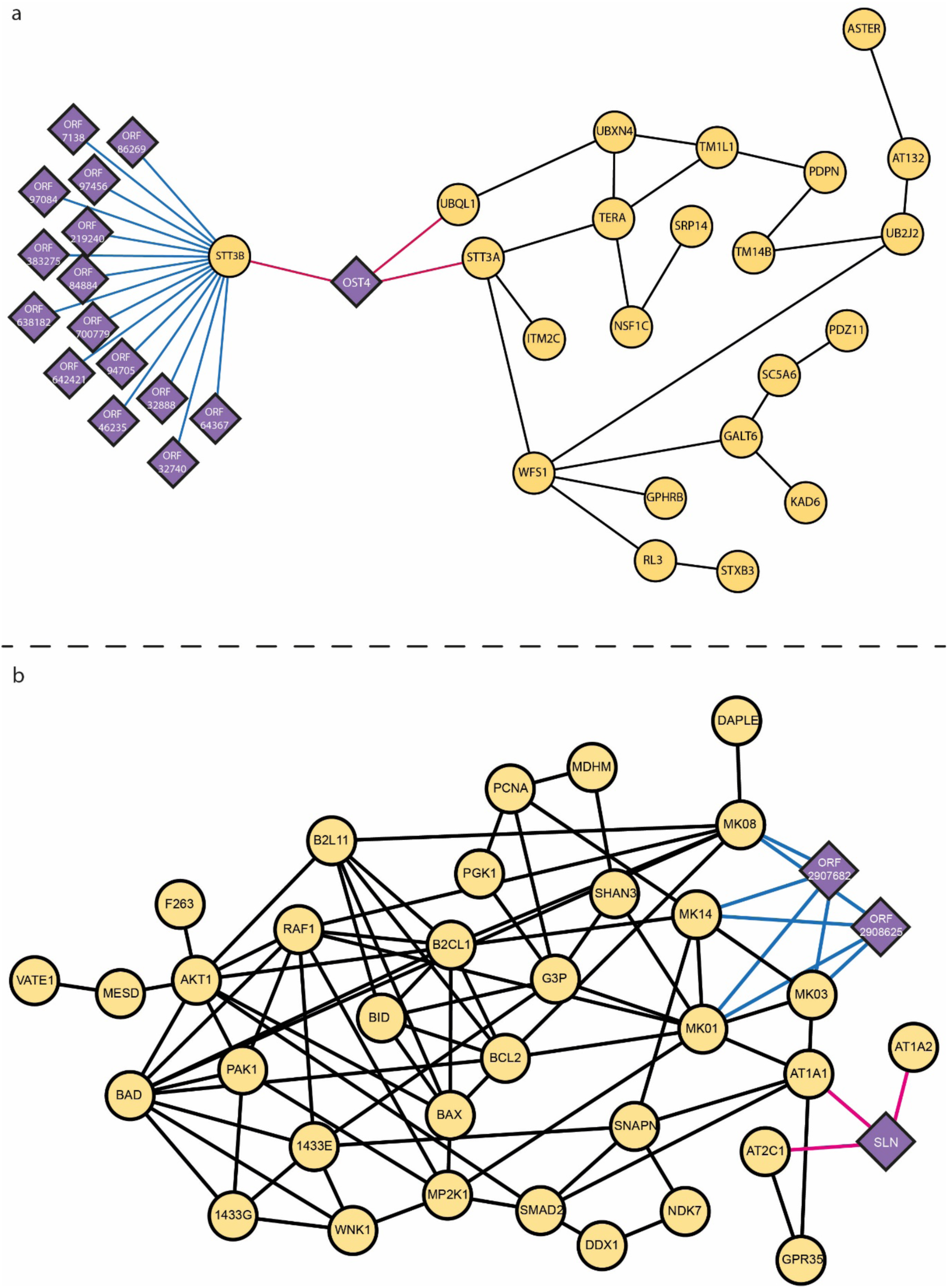
Examples of modules. a) OST4 module. b) SLN module. Purple node: sPEP. Yellow node: canonical protein. Blue edge: DMI interaction; pink edge: DDI interactions.

*SLN in skeletal muscle*: Among the 236 sPEPs identified in the skeletal muscle interactome, only one corresponds to a previously annotated protein in UniProt, sarcolipin (SLN). Current UniProt annotations describe SLN primarily through its molecular role in the regulation of ATPase activity. Consistent with the established function, we inferred direct interactions between SLN and the sarcoplasmic/endoplasmic reticulum calcium ATPases SERCA1 (ATP2A1) and SERCA2 (ATP2A2), both supported by domain–domain interaction predictions (Fig. 6b). Notably, the same interaction domain identified in SLN^67^ is also able to interact with five additional calcium-transporting ATPases (ATP2A3, ATP2B1, ATP2B2, ATP2B4, and ATP2C1) and several non-calcium ATPases, including the Na⁺/K⁺-ATPase catalytic subunits ATP1A1, ATP1A2, ATP1A4, and ATP13A3. These observations not only recover the established molecular function of SLN but also suggest that its ATPase-regulatory activity may extend beyond the SERCA pump to additional classes of P-type ATPases through a conserved interaction interface. SLN is thereby possibly involved in biological processes related to the regulation of cell communication, intracellular signal transduction, MAPK cascade activity, ERBB signaling and insulin-like growth factor receptor signaling, as suggested by the functional module annotations.

*AF1q in brain:* Among the 8014 sPEPs identified in the brain interactome, 14 correspond to previously annotated proteins in UniProtKB. One representative example is AF1Q, which is currently annotated as a cofactor of the transcription factor TCF7 and has been implicated in apoptotic signaling and positive regulation of DNA-templated transcription^68^. AF1Q is located within the same overlapping community module as TCF7, consistent with its established role as a transcriptional cofactor. Functional inference based on the organization of this interaction module, associated AF1Q with a broad spectrum of transcription-related biological processes, (e.g., “RNA polymerase II-mediated transcription”, “regulation of gene expression” and “cell differentiation”). These inferred functions substantially extend the current UniProtKB annotation while remaining fully consistent with the known biology of AF1Q as a regulator of transcriptional activity and cell differentiation in brain^68–70^.

Overall, these examples suggest that inferred annotations reflect the known functions of these sPEPs, therefore increasing confidence in our functional predictions.

## Conclusions

What are the functions of the myriad of microproteins that are detected by different meansDo they serve the same biological processes as the canonical proteinsAre they simply smaller proteins than the canonical ones that have been previously ignoredTo help answer such questions, we generated a functional landscape of the sPEPs by predicting their cellular function(s) at large-scale in three different tissues.

We chose to examine sORFs identified in RiboSeq experiments across different tissue types (and collected in the MetamORF database^27^) to detect possible tissue-specific features, therefore covering a broader functional landscape. To evaluate the protein-level evidence of our sORF dataset, we compared it to the standardized catalog of 7264 human non-canonical ORFs identified by Ribo-seq experiments^9^ proposed by the TRANSCODE consortium during the course of this work (https://transcodeconsortium.org/), a quarter of which has been detected independently by proteomic experiments^11^. Whereas the sORFs investigated in our study overlap with 2% of the catalog (144/7264), 36% of them (52/144) yielded detectable peptides. Although the overlap is low, it is significantly enriched for sORFs with proteomic evidence (Fold Change=1,5; *P*-value= 5 x 10^-4^), therefore increasing the confidence in the translation of a large part of the sORFs of our dataset. In addition, we showed that none of the tested degradation determinants indicate the sPEPs in our dataset are particularly unstable.

We predict sPEP interaction with canonical protein using mimicINT, a framework based on domain-domain and SLiM-domain interaction templates. Because SLiM-binding domains of the same family (eg. SH2, PDZ) differ in specificity^71^ and SLiMs are degenerate sequences thus prone to occur by chance, mimicINT respectively applies two filters: *(i)* a confidence score to ensure the identification of specific SLiM–domain interfaces^33,41^; *(ii)* a probability, estimated by Monte Carlo simulation, that a given motif occurs by chance in a query sequence. Applying these filters resulted in interfaces with an experimental validation rate of 70%^61^. Noticeably, we choose not using AlphaFold for interaction predictions for the following reasons: because of the short size of the sPEPs, their poor conservation, their lack of close structural homologs, and their disorder propensity, AlphaFold would not have provided reliable interaction predictions for most of the sPEPs. Indeed, not only AlphaFold performs poorly on proteins lacking related sequences and on disordered regions, but AF-Multimer predictions of domain-motif interactions are not specific when using small sequences^72^.

The analysis of the interaction interfaces among the sPEPs, SLiMs and domains, showed that sPEPs can be regulated by modifications, e.g., phosphorylation and cleavage, suggesting that they can themselves perform regulatory functions by participating in cascades of molecular events that regulate a particular biological process. Interestingly, whereas SLiM-bearing sPEPs appear to be mainly involved in signaling mechanisms in all tested tissue, domain-bearing sPEPs are devoted to tissue-specific functions. These two categories of functional sPEPs may also be distinguished by their interaction types, the first interacting transiently, whereas the latter may interact permanently^73^. We showed that most of the sPEPs interact through SLiMs, which are usually located in disordered regions, that may have a higher evolutionary rate than structured regions^74^. sPEPs having low evolutionary constraints^75^, their overall involvement in regulatory mechanisms as a group through SLiM-mediated interactions, whereas those mechanisms are usually thought to be highly specific, strongly suggest that sPEPs can represent a reservoir of functional novelty. Indeed, the fact that sPEPs could be produced by pervasive translation^11^ on the one hand, and that some of them are products of de novo genes^75^ on the other hand, suggests that sPEPs form an additional layer of complexity, acting as a raw material, malleable enough and amenable to evolutionary adaptation and selection.

## Funding sources

This work has been supported by the ”Investissements d’Avenir” French Government program managed by the French National Research Agency (ANR-16-CONV-0001) and by the Excellence Initiative of Aix-Marseille University-A*MIDEX as well as by Human Frontier Science Program grant to ARC, EBB and CB (RGP004/2023). SAC received a fellowship from the “Espoirs de la recherche” program managed by the “Fondation pour la Recherche Médicale” (FDT202106013072). Part of the bioinformatics analyses were performed on the HPC facilities of the National Network of Computing Resources of the “Institut Français de Bioinformatique” (IFB), funded by the “Programme d’Investissements d’Avenir” (PIA), grant Agence Nationale de la Recherche (ANR-11-INBS-0013).

## Acknowledgements

“Centre de Calcul Intensif d’Aix-Marseille” is acknowledged for granting access to its high-performance computing resources. Hugo Bonnefous and Jean-Christophe Mouren are particularly thanked for their help in statistics and code debugging.

## Author contributions

M.S. and S.A.C. developed the analysis pipeline, performed the data analyses and drafted the manuscript. L.S. & L.S. contributed to code development and interactome building. P.P. participated to the supervision of the start of the project. A.Z. contributed to code implementation, computational tool usage, and conceptualization. A-R.C. and E.B.B reviewed and edited the manuscript. C.B. conceptualized, conceived, supervised, and acquired funds for the project. M.S. and C.B. wrote the manuscript based on a first draft from

S.A.C. and C.B., with input from all authors.

## Data availability

Data supporting the findings of this study are available in a Zenodo repository (https://doi.org/10.5281/zenodo.11191677) organized by analysis blocks in accordance with the structure of the paper. All datasets used in this study are publicly available or have been deposited in appropriate repositories.

## Code availability

We deposited codes and bioinformatics environments in GitHub at (https://github.com/TAGC-NetworkBiology/InteractORF) and in Zenodo (https://doi.org/10.5281/zenodo.22689845). Both data and codes are publicly available for the replication of the whole study.

## Competing interests

All authors declare no competing interest.

## Notes

### Competing Interest Statement

The authors have declared no competing interest.

### Summary of Updates

The analyses have been extended to two more tissues and considerably improved.

